# Serotonergic and catecholaminergic (dopaminergic) oscillations in the reproductive regulation of Japanese quail

**DOI:** 10.1101/345017

**Authors:** Suneeta Yadav, Chandra Mohini Chaturvedi

**Affiliations:** Department of Zoology, Banaras Hindu University, Varanasi 221 005, India (Email-,)

**Author notes:** Corresponding Author:- Prof. (Mrs.) Chandra Mohini Chaturvedi, Department of Zoology, Banaras Hindu University, Varanasi-221 005, U.P., INDIA, 9415819743.

**Keywords:** 5-HTP, L-DOPA, Specific temporal phase relation, Serotonergic and dopaminergic oscillations

## Abstract

Specific temporal phase relation of serotonergic and dopaminergic oscillations alters reproductive responses in many species. Aim of the study was to confirm whether effect of serotonergic drug (5-HTP) and dopaminergic drug (L-DOPA) is due to their conversion into serotonin and dopamine respectively or other products. For this study, PCPA (p-chlorophenylalanine, a long lasting inhibitor of serotonin synthesis), DDC (Diethyldithiocarbamate, which inhibits biosynthesis of nor-adrenaline), α-MT (Methyl-p-tyrosine, an inhibitor for the conversion of tyrosine to DOPA) and DOPS (Dihydroxyphenylserine, a specific precursor for noradrenaline) were used in different groups in addition to 5-HTP and L-DOPA given at specific time interval. Reproductive responses monitored at 10 weeks post treatment indicate that gonadal activity was significantly low in HTP:DOPA (8-hr quail), HTP+PCPA:DOPA and HTP:DOPA+DDC quail compare to control (S:S). However, gonadal activity of HTP:S(HTP control), S:DOPA(DOPA control) and HTP: α-MT+DOPS was not different from S:S control and remained in active condition. These findings indicate that it is not the dose of neurotransmitter precursor drugs (5-HTP and L-DOPA) and the neurotransmitters (serotonin and dopamine itself) that cause the effect, instead it is the function of interval between the drug administration which induces or entrains specific phase relation between serotonergic and dopaminergic oscillations. Further, gonadal suppression observed in HTP:DOPA, HTP+PCPA:DOPA and HTP:DOPA+DDC group three groups is not due to injection of 5-HTP or L-DOPA (alone) but due to conversion of administered 5-HTP into serotonin and conversion of L-DOPA (administered) into dopamine; not due to their further conversion into catecholamines other than dopamine i.e. noradrenaline or adrenaline.

## Introduction

The studies on neurotransmitters proves that the brain levels of nor-epinephrine (Owasoyo and Walker, 1980), dopamine (Owasoyo *et al*., 1979), acetylcholine (Saito, 1971) as well as pineal melatonin (Lynch, 1971) were highest during the dark phase of the light:dark cycle and lowest during the light phase where as serotonin (Quay, 1968; Owasoyo and Walker, 1980) shows an opposite circadian pattern with peak levels occurring during light phase of the light:dark cycle. It has also been shown that brain serotonin and other neurotransmitters (dopamine) exhibit circadian variation in different brain areas including suprachiasmatic nuclei (SCN) (Philo *et al*., 1977; Héry *et al*., 1981; Wilson and Meier, 1987; 1988). Experimentally it has been demonstrated that the circadian rhythms exist in hormones as well as neurotransmitters viz. 5-HT, NE and DA (Manshardt and Wurtman, 1968; Wilson and Meier, 1987; Forsling, 2000; Tiwari *et al*., 2006). A number of studies also indicate the circadian release of 5-HT in blood and pineal of mammals, birds and fish (Reis and Wurtman, 1968; Reis *et al*., 1969). Thus, there appears to be considerable synchrony throughout the brain with regards to the rhythms of neurotransmitters content and activity (Reis *et al*., 1969; Le Bras, 1984; Khan and Joy, 1990).

It has been well established from several reports that administration of 5-HTP and L-DOPA at the interval of 12 hrs stimulates gonadal growth and body weight gain whereas administration of these drugs at the interval of 8 hrs results into opposite effect i.e. gonadal suppression in several avian (Red headed bunting-Chaturvedi and Bhatt, 1990; Bhatt and Chaturvedi, 1992b; Phillips and Chaturvedi, 1992; Lal munia- Chaturvedi et al., 1994; Japanese quail- Chaturvedi et al., 1991; Bhatt and Chaturvedi, 1992a; Phillips and Chaturvedi, 1995; Bhatt and Chaturvedi, 1998; Tiwari and Chaturvedi, 2003; Chaturvedi et al., 2006; Kumar and Chaturvedi, 2008; Indian weaver bird- Chaturvedi et al., 1997; Spotted munia- Chaturvedi and Prasad, 1991; Prasad and Chaturvedi, 1992a, 1992b, 1992c, 2003) and mammalian species whether seasonally breeding (Syrian hamster- Wilson and Meier, 1989; Indian palm squirrel- Chaturvedi and Jaiwal, 1990; Jaiwal and Chaturvedi, 1991; Chaturvedi and Singh, 1992) or continuous breeder (laboratory mouse, *Mus musculus*- Sethi and Chaturvedi, 2009; Sethi et al, 2010).

In some of these studies, instead of 5-HTP and L-DOPA given at specific time interval, each drug was given in combination with saline (5-HTP & saline or saline & L-DOPA) (Prasad and Chaturvedi, 2003). Since these combinations did not produce any significant effect or long lasting effect, it was suggested that it is not the effect of either serotonin or dopamine alone, but actually the interval between the administrations of two drugs is important to mimic the seasonal gonadal condition. It was presumed and later on also proved experimentally that, these timed injection of serotonin and dopamine precursor drugs given for a period of 11-13 days may entrain circadian serotonergic and dopaminergic oscillation respectively. Moreover, the two precursor drugs given at different time interval will induce different phase relationship or phase angle between the two oscillations and different physiology (Yadav and Chaturvedi, 2014, 2015). Obviously, in nature also during sexually active and inactive condition these phase relations vary the underlying basis of our hypothesis and all the experimental studies in this direction (Wilson and Meier, 1987, 1988, 1989; Tiwari and Chaturvedi, 2003; Tiwari et al., 2006). In view of fact that neurotransmitters serotonins and dopamine do not cross the blood brain barrier but their precursor do cross (Bianchine, 1980) in all these studies 5-HTP and L-DOPA were used as the precursor of serotonin and dopamine respectively instead of neurotransmitter itself.

In case of Spotted munia, *Lonchura punctulata*, it has been proved that these effects are due to temporal phase relationship of circadian neural oscillations and not due to serotonin and dopamine alone (Prasad and Chaturvedi, 2003). L-DOPA not only gets converted into dopamine in the presence of enzyme tyrosine hydroxylase but the next biosynthetic product is the noradrenaline (by enzyme dopamine hydroxylase) and adrenaline (by enzyme N-methyl transferase) respectively. Further, tryptophan converts into 5-HTP (by tryptophan hydroxylase) and 5-HTP gets converted into serotonin (by enzyme aromatic amino acid decarboxylase) and next product is melatonin in certain tissues. This experiment was conducted to confirm that effects observed in the earlier studied are due to conversion of 5-HTP into serotonin and that of L-DOPA into dopamine only and not their next biosynthetic products. Hence agonist and antagonists of different enzymes were used an addition to 5-HTP and L-DOPA given at specific time interval to study the reproductive response of Japanese quail. The biosynthetic pathway of neurotransmitter serotonin and dopamine, conversion of dopamine into other catecholamines and agonists and antagonists of these biosynthetic products are described below in details along with the line diagram.

1. PCPA (p-chlorophenylalanine), a long lasting inhibitor of serotonin synthesis (Koe and Weiseman, 1966). It blocks the conversion of tryptophan into 5-HTP by inactivating the enzyme tryptophan hydroxylase. It does not affects the conversion of 5- HTP into serotonin.
2. DDC (Diethyldithiocarbamate), which inhibits the conversion of Dopamine to nor-adrenaline by inducing negative effect on the enzyme Dopamine ß-hydroxylase, essential for this biosynthesis (Crevelling et al., 1968).
3. α-MT (Methyl-p-tyrosine). An inhibitor for the conversion of tyrosine to DOPA by affecting the enzyme tyrosine hydroxylase.
4. DOPS (Dihydroxyphenylserine), a specific precursor for noradrenaline. So it enhances the synthesis of noradrenaline.

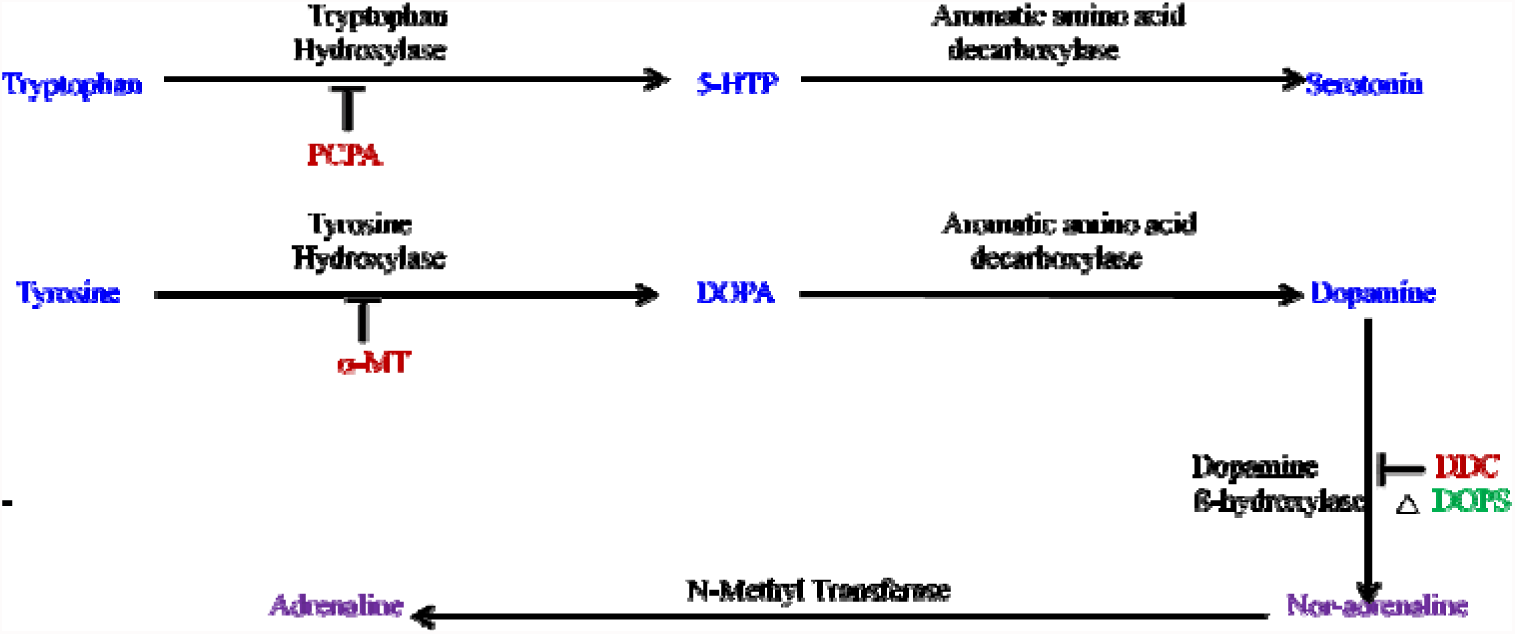

The specific aim of the present study was to prove that stimulation (due to 12 hr phase relation) or regression (due to 8 hr phase relation) of gonadal development is due to specific temporal phase relation of serotonergic and dopaminergic activities or oscillations and not due to i) serotonin/5-HTP or dopamine/L-DOPA alone and ii) that L-DOPA was effective when converted into dopamine and not into noradrenaline or adrenaline. For the above mentioned objectives, some agonist/antagonists of serotonin (PCPA) and Dopamine (α-MT, DDC, DOPS) were used in combination with these neurotransmitters (serotonin and dopamine) precursors (5-HTP and L-DOPA).

## Materials and methods

Three week old male Japanese quail purchased from Chuck Gazaria farm, Lucknow, were divided into 7 groups each having 8 birds.

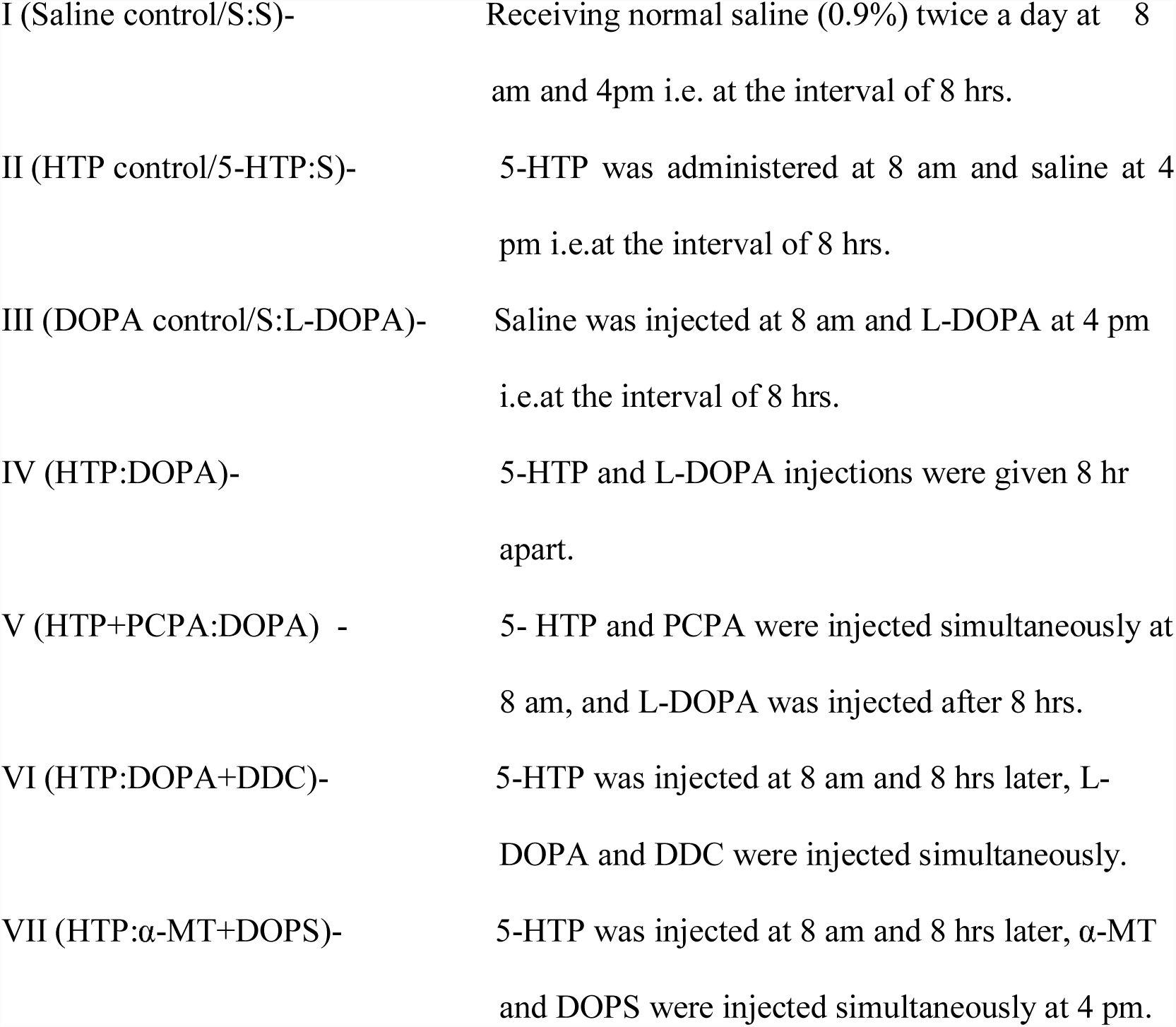

All the above injections were given intraperitoneally in 100 µl saline at 8 am and 4 pm so as to establish 8-hr interval/phase relationship between the two injections or set of injections. The dose of 5-HTP and L-DOPA was 5mg/100g body weight per day for 13 days. The doses of antagonists were as follows- PCPA (2mg/100 g body wt.) DDC (10 mg/100g body wt.), α-MT (1 mg/100g body wt.) and agonist of noradrenaline DOPS was (10mg/100 g body wt.). During the treatment of 13 days, all the birds were kept in dim continuous light (2 lux) and after the treatment period, birds of all the groups were transferred to long day condition (LD 16:8).

The length and width of the cloacal gland was measured *in situ* with dial calipers weekly; before, during and after the treatment upto the termination of study and cloacal gland volume was calculated in cm^3^ (Jaiwal and Chaturvedi, 1991; Chaturvedi *et al*., 1993). At the terminations of study i.e. 10 weeks post treatment (at the age of 15 weeks), blood was collected from the alar vein/wing vein into a heparinized tube and centrifuged at 4000 rpm for 20 min at 4°C to separate plasma and stored at -20° for hormonal assays to be performed later. Plasma testosterone level was measured using EIA kit (DSI s.r.l., Italy) following manufacturer’s protocol. The antiserum used in the assay was 100% specific for testosterone (cross reactivity/specificity with testosterone was 100%); the cross reactivity of the assay was 0.056% with progesterone, 0.004% with cortisol, 0.005% with estradiol, 4.8% with dihydrotestosterone, 3.6% with androstenedione, 0.048% with androsterone, 0.004% with cortisone, 0.002% with estriol and 0.007% with estrone. The analytical sensitivity of the assay was 0.0576 ng/ml. The intra-assay coefficient of variation (CV) is 5.6% whereas inter-assay CV is 7.1%. Accuracy for this assay was 99%. Thereafter, birds from each group were weighed, deeply anaesthesized with thiopentone and then dissected to collect tissues so as to process for measuring testicular volume (cm^3^) and for calculating gonadosomatic index-GSI (in gram testes/100 gram body weight). Left testis of each quail after measuring its length and width *in situ* for calculating testicular volume was fixed in Zamboni’s solution and processed for routine histological study and measurement of seminiferous tubule diameter. For histology, twenty-four hours after fixation, the testes were dehydrated in an ascending series of alcohol, treated with xylene and then embedded in paraffin wax. The 6-μm thick sections were cut by a Weswox rotary microtome (Western Electric and Scientific Works, Ambala Cantt, India), and stained with hematoxylin-eosin. Histological sections were viewed under a microscope (Axioskop 2 Plus; Carl Zeiss AG, Oberkochen, Germany) and images were captured with a digital camera. The diameter of the seminiferous tubules was determined in 10 sections from each testis by using the occulometer and micrometer.

All the numerical data (cloacal gland and testicular volume, GSI, seminiferous tubule diameter and plasma testosterone concentration) were analyzed by one-way analysis of variance (ANOVA), followed by post hoc Dunnett test for the comparison of group means. Significance was calculated at the level of p<0.05.

## Results

Cloacal gland volume of quail of all the groups maintained under LD 16:8 remained suppressed during the period of treatment (until 5 weeks of age). Thereafter a sharp increase until 9 weeks of age followed by maintenance of plateau at increased level was observed in saline: saline (S:S) i.e. saline control, HTP: saline i.e. HTP control, saline: DOPA i.e. DOPA control and HTP: α-MT+DOPS group quail. However, those of HTP:DOPA, HTP+PCPA:DOPA and HTP:DOPA+DDC remained at significantly low level throughout the period of study compare to control (saline:saline) (Fig. 1). At the termination of study, testicular volume, GSI and plasma testosterone level of HTP:DOPA, HTP+PCPA:DOPA and HTP:DOPA+DDC group quail were significantly lower in comparison to saline control (S:S) but these parameters of other 3 groups were not different from saline control (Fig. 2, 3 and 4).

**Fig. 1.**
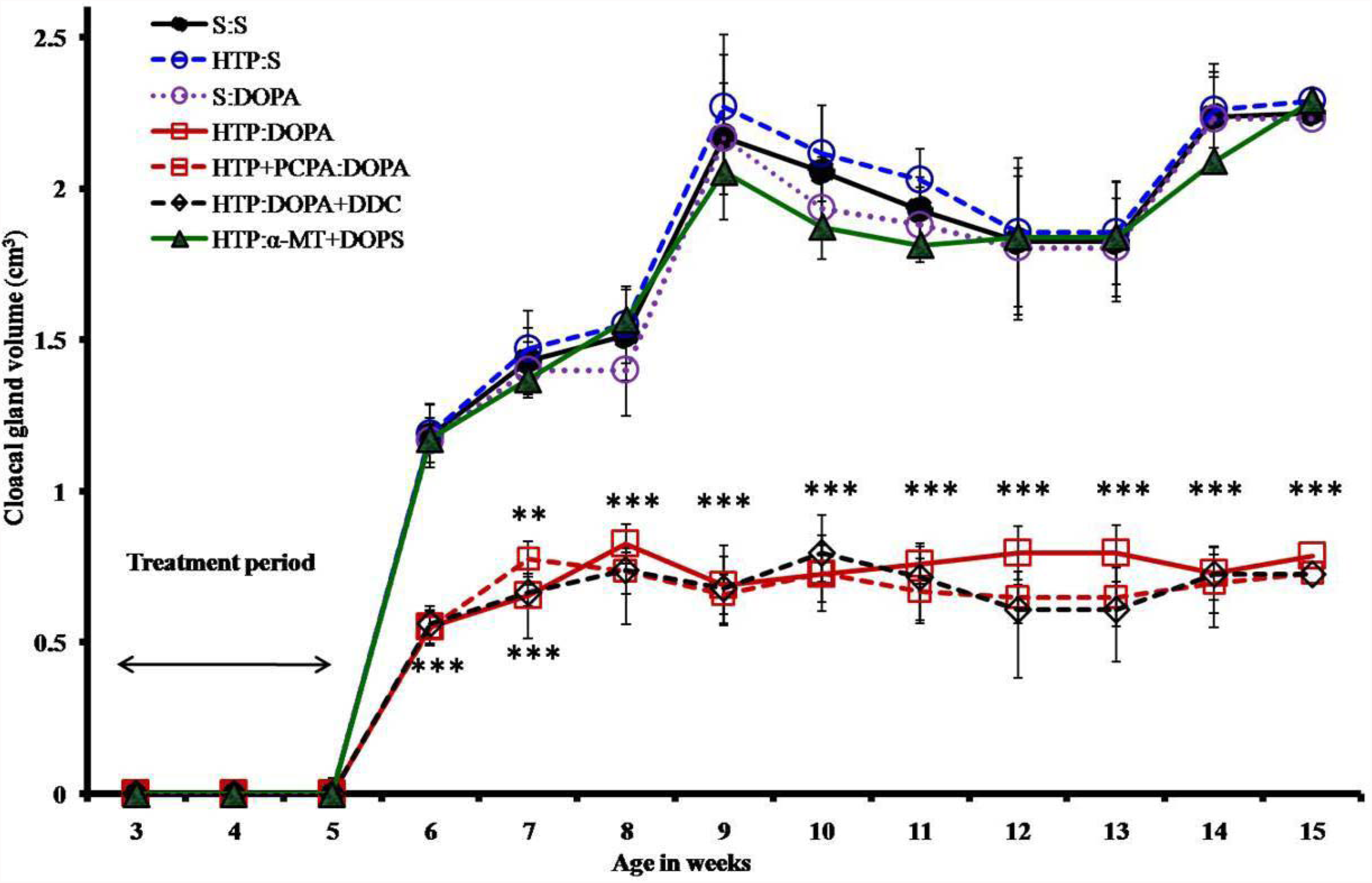
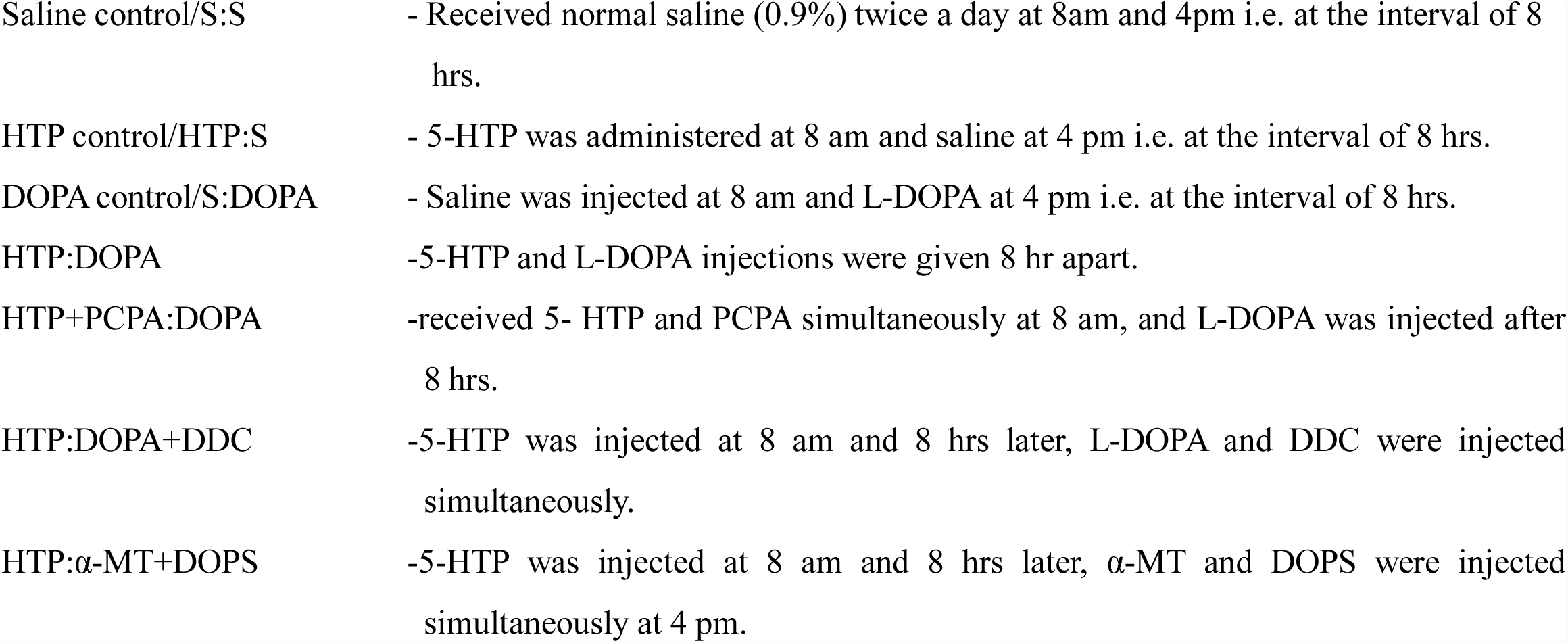
Cloacal gland responses of Japanese quail receiving 5-HTP and L-DOPA in combination with saline and agonist and/or antagonist of serotonin and catecholamines 8 hrs apart. Data is presented as means ± SEM. **p<0.01, ***p<0.001, Significance of difference from the control group. Group details:

**Fig. 2.**
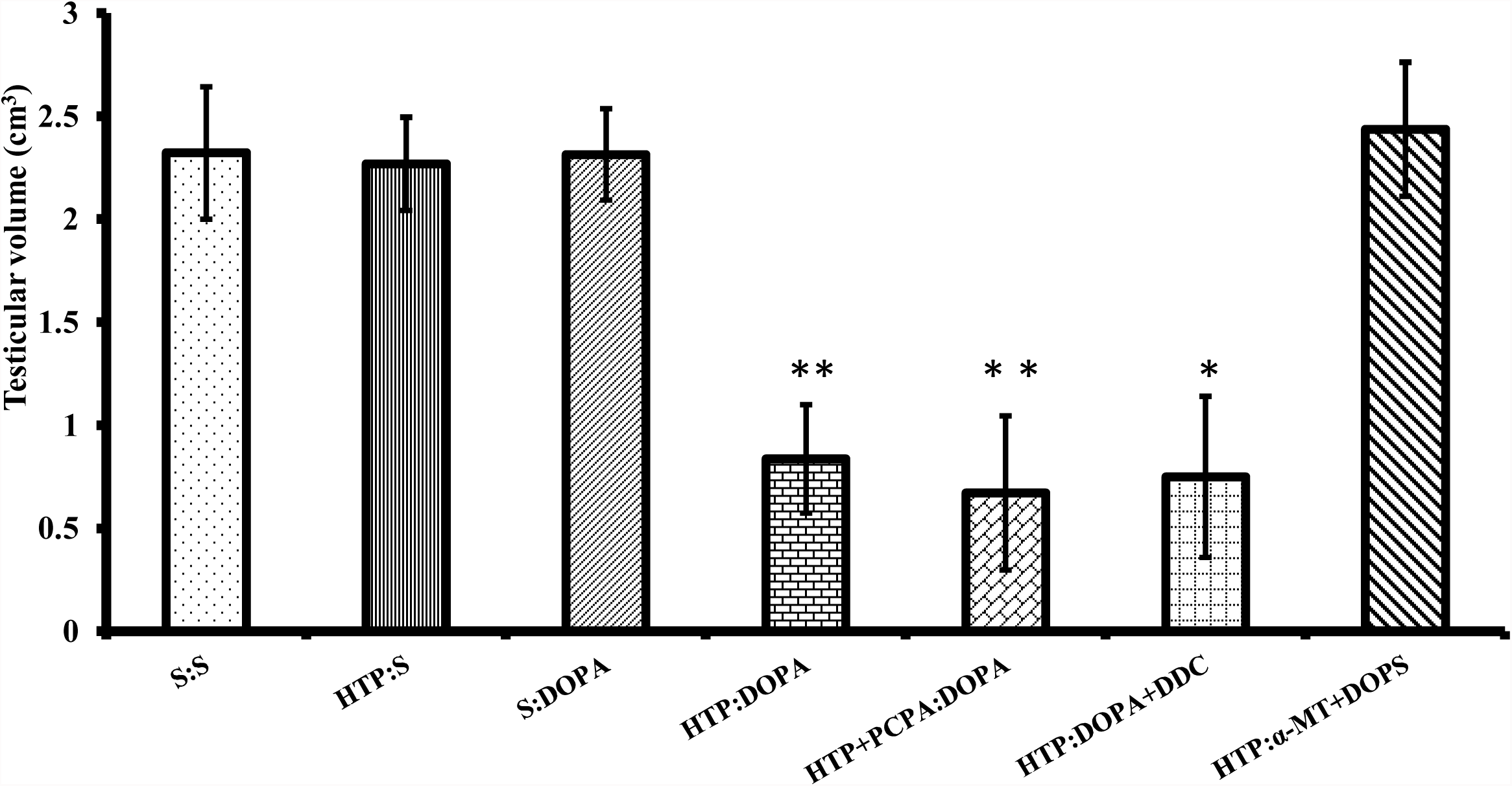
Testicular volume of Japanese quail receiving 5-HTP and L-DOPA in combination with saline and agonist and/or antagonist of serotonin and catecholamine 8 hrs apart. For group details, see Fig. 1. Data is presented as mean ± SEM. *p<0.05, **p<0.01; Significance of difference from the control (S:S) group.

**Fig. 3.**
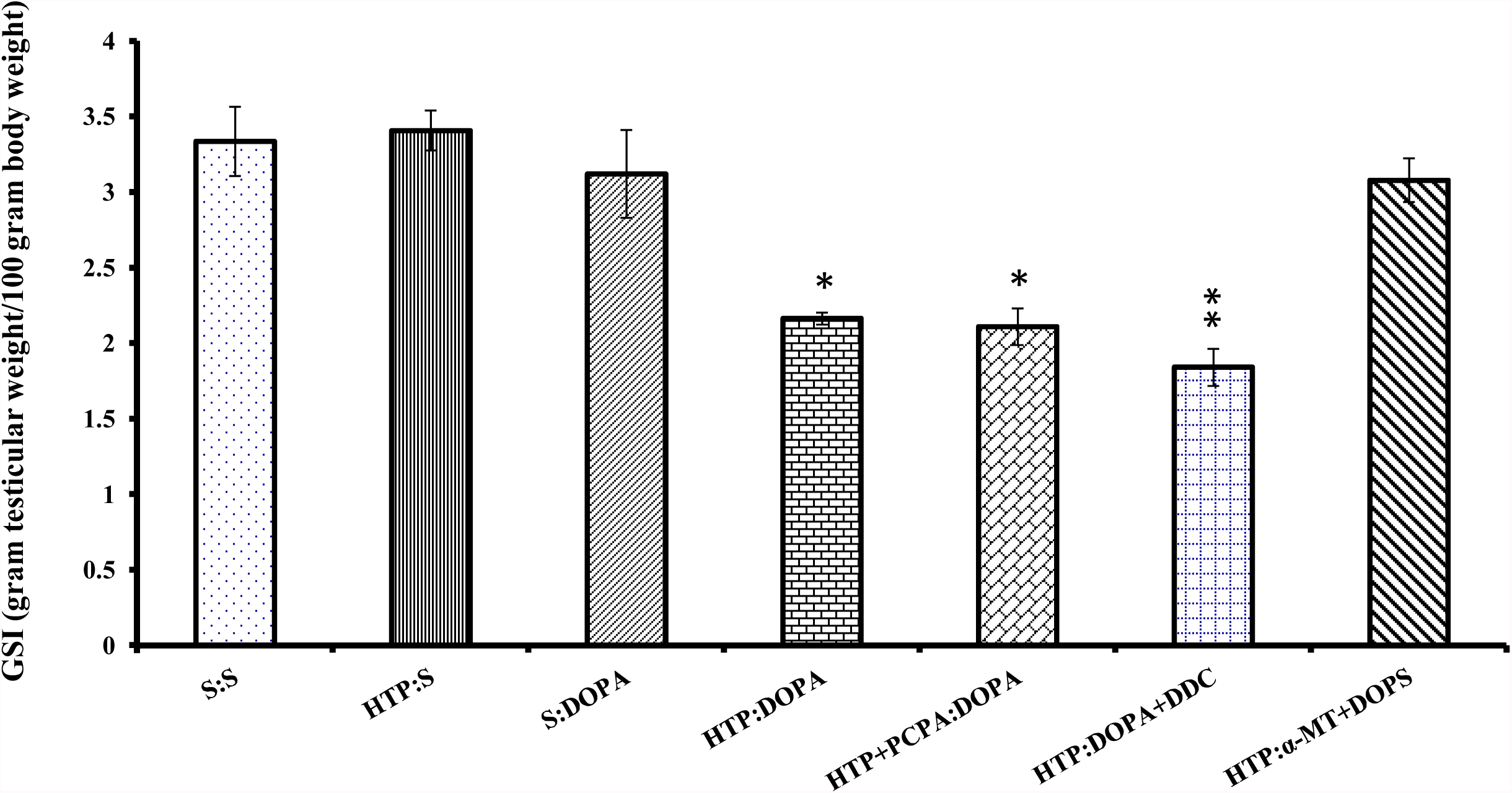
GSI (Gonadosomatic index) of Japanese quail receiving 5-HTP and L-DOPA in combination with saline and agonist and/or antagonist of serotonin and catecholamine 8 hrs apart. For group details, see Fig. 1. Data is presented as mean ± SEM. *p<0.05, **p<0.01; significance of difference from the control group

**Fig. 4.**
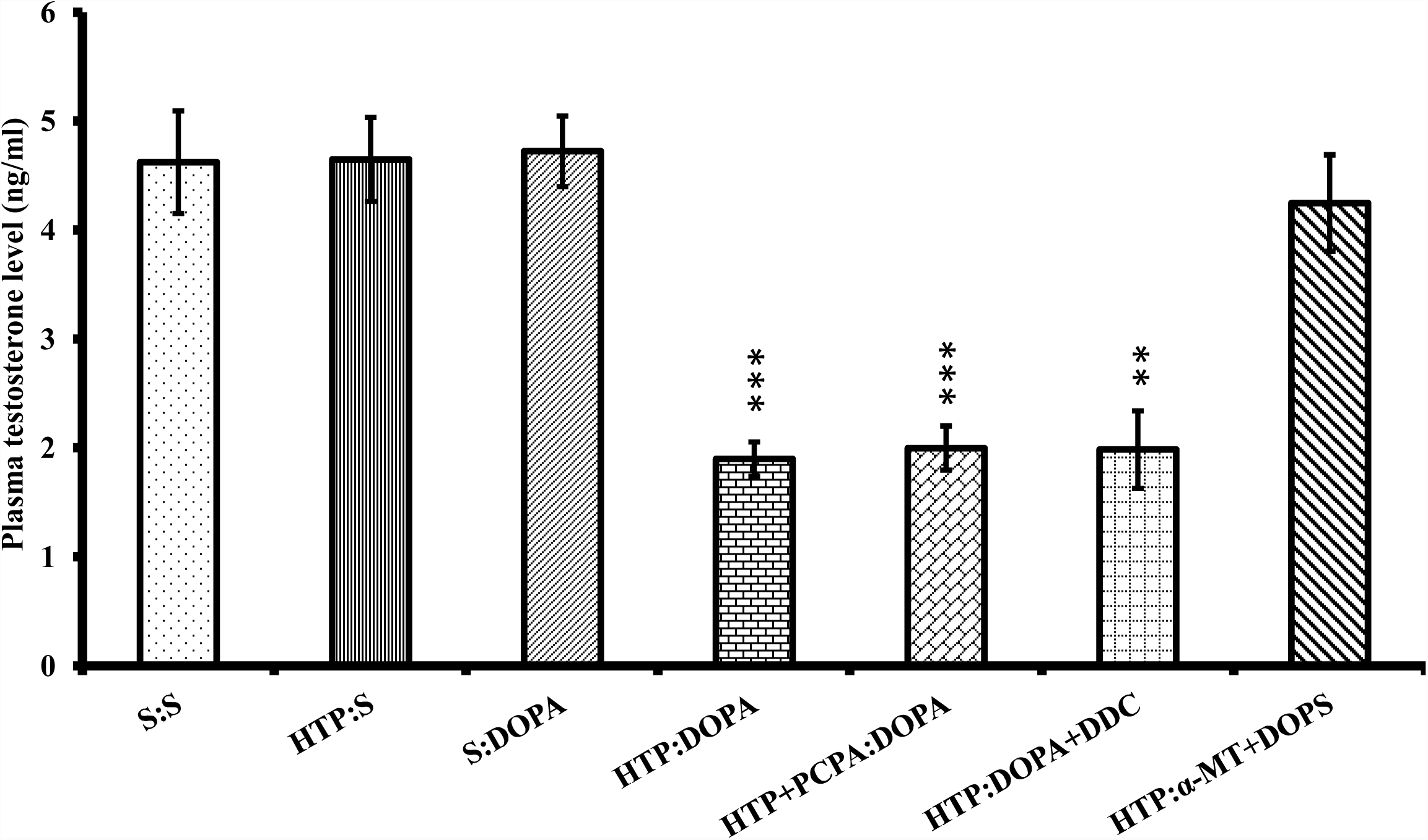
Plasma testosterone level of Japanese quail receiving 5-HTP and L-DOPA in combination with saline and agonist and/or antagonist of serotonin and catecholamine 8 hrs apart. **p<0.01, ***p<0.001; Significance of difference from the control group.

Histologically, the transverse section of testis of HTP:DOPA, HTP+PCPA:L-DOPA and HTP:DOPA+DDC 8-hr quail group had smaller seminiferous tubules (Fig. 5) with decreased spermatogenesis along with vacuolation in HTP:DOPA and HTP:DOPA+DDC quail and atrophic changes with complete degeneration of spermatogenenic activity was noted in HTP+PCPA:DOPA quail testis unlike full breeding condition in control (S:S). The interstitial spaces of the seminiferous tubules of these quail testes were reduced and no Leydig cells were evident in triangular spaces. Further, similar to the enlarged seminiferous tubules of saline control quail testis having all the stages of spermatogenesis and bunches of spermatozoa in the lumen, the testis of HTP:S, S:DOPA and HTP: α-MT+DOPS also exhibited full breeding condition (Fig. 5 and 6).

**Fig. 5.**
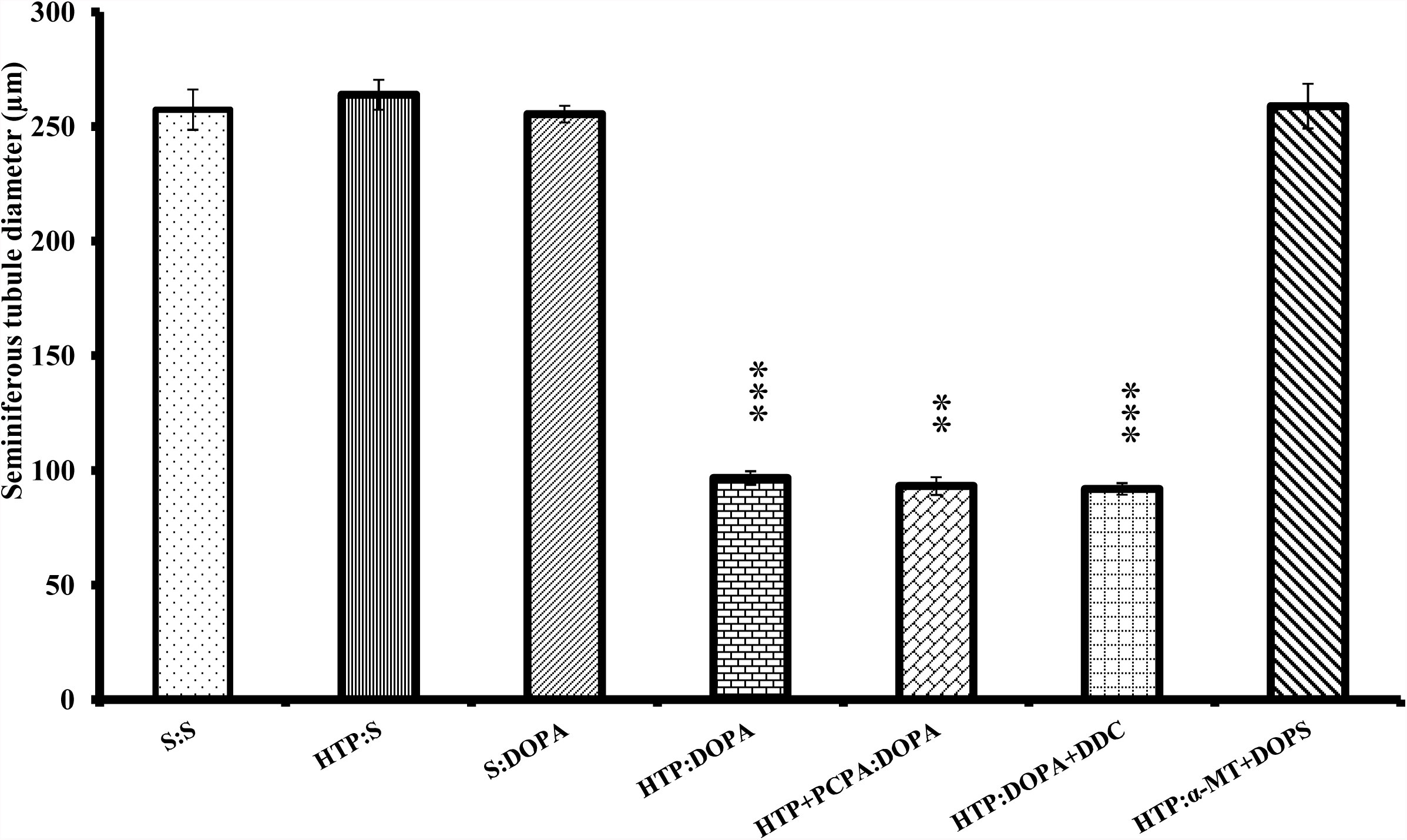
Seminiferous tubule diameter of Japanese quail receiving 5-HTP and L-DOPA in combination with saline and agonist and/or antagonist of serotonin and catecholamine 8 hrs apart. **p<0.01, ***p<0.001; Significance of difference from the control group.

**Fig. 6.**
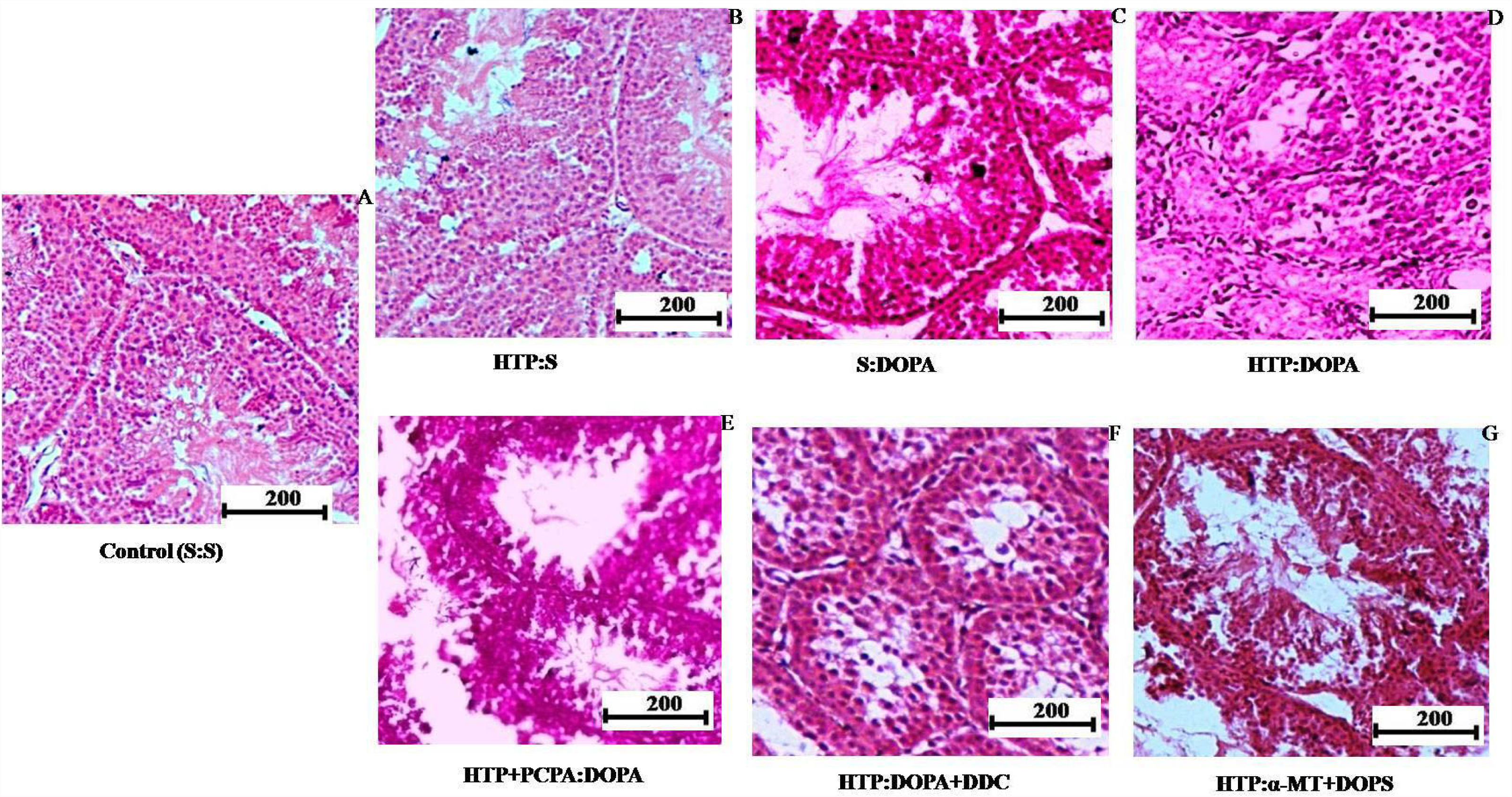
T.S. of testis of Japanese quail receiving 5-HTP and L-DOPA in combination with saline and agonist and/or antagonist of serotonin and catecholamines 8 hrs apart. A. Control (S: S)- Note full breeding condition of testis having enlarged seminiferous tubules with all the stages of spermatogenesis and spermatozoa in the lumen. B. HTP control (HTP: S)- showing full breeding condition as in S:S quail testis. C. DOPA control (S:DOPA)- showing full breeding condition as in S:S quail testis. D. HTP:DOPA- Note non breeding condition with smaller seminiferous tubules containing only inactive spermatogonial cells and some vacuolation and debrises in the lumen of few tubules E. HTP+PCPA:DOPA- Note smaller seminiferous tubules with suppression of stages of spermatogenic activity and emply lumen or some debrises in lumen . F. HTTP:DOPA+DDC- Note non-breeding and spermatogenetically inactive condition having smaller seminiferous tubules containing inactive spermatogonial cells and vacuolation. G. HTP: α-MT+DOPS group receiving 5-HTP and α**-**MT +DOPS at the interval of 8 hr. This section shows normal breeding condition like control.

## Discussion

Intraperitoneal injections of 5-HTP and L-DOPA (HTP:DOPA) given 8 hrs apart i.e. 8-hr phase relation of serotonergic and dopaminergic oscillations induced by their precursor injections suppressed gonadal activity significantly compared to saline treated control (S:S). On the other hand, reproductive parameters of quail receiving saline 8 hrs after 5-HTP (HTP:S) or 8 hrs before DOPA (S:DOPA) serving as HTP control and DOPA control respectively, did not differ from saline control (S:S). This indicates **t**hat suppressive effect of 5-HTP and L-DOPA when given 8 hrs apart (8-hr relation) was neither due to 5-HTP nor L-DOPA alone but is actually the outcome of the interval between the administration of serotonergic and dopaminergic precursor drugs inducing 8-hr temporal relationship between the two neural oscillations. In another group of birds, 5-HTP and PCPA was injected simultaneously followed by L-DOPA injection at interval 8 hr. PCPA blocks the conversion of tryptophan into 5-HTP by inactivating the enzyme tryptophan hydroxylase but it does not affect the conversion of 5-HTP into serotonin. Hence exogenous administration of PCPA blocked the conversion of endogenous tryptophan into 5-HTP, but exogenous 5-HTP was still converted into serotonin. Hence in fact, theoretically 8-hr phase relation of HTP: DOPA should not be different from HTP+PCPA:DOPA and accordingly gonadal response of these two groups are also similar i.e. gonado-suppressive.

In case of HTP:DOPA+DDC quail also, gonadal suppression was observed similar to those quail receiving only two drugs HTP:DOPA 8 hrs apart. The injection of DOPA (dopamine precursor) increases the synthesis of dopamine (DA) and enzyme dopamine ß-hydroxylase (DBH) converts DA into noradrenaline (NA). The drug DDC selectively inhibits NA synthesis because it inhibits the enzyme DBH therefore inhibiting the synthesis of NA. Hence simultaneous injection of DOPA and DDC is expected to stimulate dopamine synthesis but inhibits synthesis of NA. Obviously effect of this combination of drugs (HTP:DOPA+DDC) was also similar to that of HTP: DOPA combination indicating that inhibitory effect of 8-hr temporal relation of precursor drugs was only due to their conversion into serotonin and dopamine and not any further conversion into noradrenaline.

In the HTP: α-MT+DOPS group, quail received α-MT and DOPS simultaneously 8 hrs after 5-HTP administration and the effect was not different from control. The drug α-MT inhibits the conversion of the amino acid tyrosine to DOPA, whereas the drug DOPS is a specific precursor for NA. Hence, simultaneous injections these two drugs (α-MT i.e. antagonist of DOPA synthesis and DOPS i.e. agonist of NA) should selectively inhibit synthesis of DOPA (and hence also the dopamine synthesis) and increase NA synthesis respectively. Because in this treatment of HTP:α-MT+DOPS, there is no exogenous DOPA (precursor of dopamine) and internal source of DOPA is also blocked by α-MT, the DA synthesis is expected to be blocked completely but NA is still available synthesized from its selective precursor DOPS. Although natural precursor of adrenaline/NA i.e. dopamine is not available (as endogenous dopamine synthesis have been blocked by α-MT, and exogenous DOPA is not available), but NA will be still available because of injecting its selective precursor DOPS. Thus this combination of drugs is equivalent to 8-hr relationship between 5-HTP/serotonin and NA (noradrenaline) and not the 5-HTP/ serotonin and L-DOPA /dopamine. But, in terms of gonadal response, effect is similar to HTP control which in turn is similar to saline control.

These various combinations of serotonergic and nor adrenergic drugs instead of classical serotonin: dopamine combination indicates that reproductive effect of specific phase relation of neural oscillations occurs due to serotonin and dopamine; and not due to serotonin and noradrenaline (Fig. 1 and 6). These experimental findings following the use of various agonists and antagonist of monoamine (serotonin) and catecholamine (dopamine and noradrenaline) suggest that reproductive effect of 5-HTP and L-DOPA given 8 hrs apart apparently result from their conversion into serotonin and dopamine and not due to other monoamine/catecholamine. Selective potentiation of dopaminergic activity by DOPA and DDC injections did suppress reproductive development whereas specific inhibition of dopaminergic activity by α-MT and stimulation of noradrenaline by DOPS injection did not. Therefore, HTP/ /serotonergic or L-DOPA/dopaminergic activity independently is unable to induce any change in gonadal activity which is actually the effect/function of interval between the administration of neurotransmitter precursor drugs 5-HTP and L- DOPA. Moreover, it is not the dose of neurotransmitter precursor drugs (5-HTP and L-DOPA) that cause the effect, instead it is the specific time interval between the drug administrations, otherwise the effects could have been similar in all cases and not the one observed in all the reports consistently. Invariably in many species, 8-hr phase relation is gonado-suppressive, 12-hr is gonado-stimulatory and other relations are ineffective (Chaturvedi and Bhatt, 1990; Chaturvedi and Yadav, 2013).

Based on these findings it is restated that effects observed in the experiments of present study, are due to specific phase relation of serotonergic and dopaminergic oscillations induced by systemic injections of 5-HTP and L- DOPA given at different interval. Present findings also suggest that the observed effect are due to conversion of DOPA into dopamine and not into the next biosynthetic product (NA).

## Acknowledgements

Junior research fellowship to Suneeta Yadav from University Grants Commission (BSR-RFSMS), New Delhi and Research projects to CMC from Department of science and Technology, New Delhi, India (SR/50/AS-14/2012) is gratefully acknowledged.

